# Creation of a novel trigeminal tractography atlas for automated trigeminal nerve identification

**DOI:** 10.1101/2020.01.15.904938

**Authors:** Fan Zhang, Guoqiang Xie, Laura Leung, Michael A. Mooney, Lorenz Epprecht, Isaiah Norton, Yogesh Rathi, Ron Kikinis, Ossama Al-Mefty, Nikos Makris, Alexandra J. Golby, Lauren J. O’Donnell

## Abstract

Diffusion MRI (dMRI) tractography has been successfully used to study the trigeminal nerves (TGNs) in many clinical and research applications. Currently, identification of the TGN in tractography data requires expert nerve selection using manually drawn regions of interest (ROIs), which is prone to inter-observer variability, time-consuming and carries high clinical and labor costs. To overcome these issues, we propose to create a novel anatomically curated TGN tractography atlas that enables automated identification of the TGN from dMRI tractography. In this paper, we first illustrate the creation of a trigeminal tractography atlas. Leveraging a well-established computational pipeline and expert neuroanatomical knowledge, we generate a data-driven TGN fiber clustering atlas using tractography data from 50 subjects from the Human Connectome Project. Then, we demonstrate the application of the proposed atlas for automated TGN identification in new subjects, without relying on expert ROI placement. Quantitative and visual experiments are performed with comparison to expert TGN identification using dMRI data from two different acquisition sites. We show highly comparable results between the automatically and manually identified TGNs in terms of spatial overlap and visualization, while our proposed method has several advantages. First, our method performs automated TGN identification, and thus it provides an efficient tool to reduce expert labor costs and inter-operator bias relative to expert manual selection. Second, our method is robust to potential imaging artifacts and/or noise that can prevent successful manual ROI placement for TGN selection and hence yields a higher successful TGN identification rate.

## 1. Introduction

The trigeminal nerve (TGN) is the largest and most complex of the 12 pairs of cranial nerves in the brain. It includes multiple segments, including brainstem, cisternal, Meckel’s cave and peripheral branches (see Figure 1 for an anatomical overview) (Go et al., 2001; Joo et al., 2014). It supplies sensation to the skin in the face, the ear, the mucous membranes orally and endonasally as well as motor innervation to the muscles of mastication. The TGN has been shown to be affected in many diseases such as trigeminal neuralgia (Jannetta, 1967), multiple sclerosis (Love & Coakham, 2001; Yadav et al., 2017), local ischemia (Balestrino & Leandri, 1997; Delitala et al., 1999; Golby et al., 1998) and brain cancer (Timothee Jacquesson et al., 2019). Many research studies have also suggested that the identification of TGN is important for understanding and/or potential treatment of various neurological disorders such as major depressive disorder (Schrader et al., 2011), attention-deficit/hyperactivity disorder (McGough et al., 2015), and Parkinson’s disease (Barz et al., 1997).

**Figure 1.**
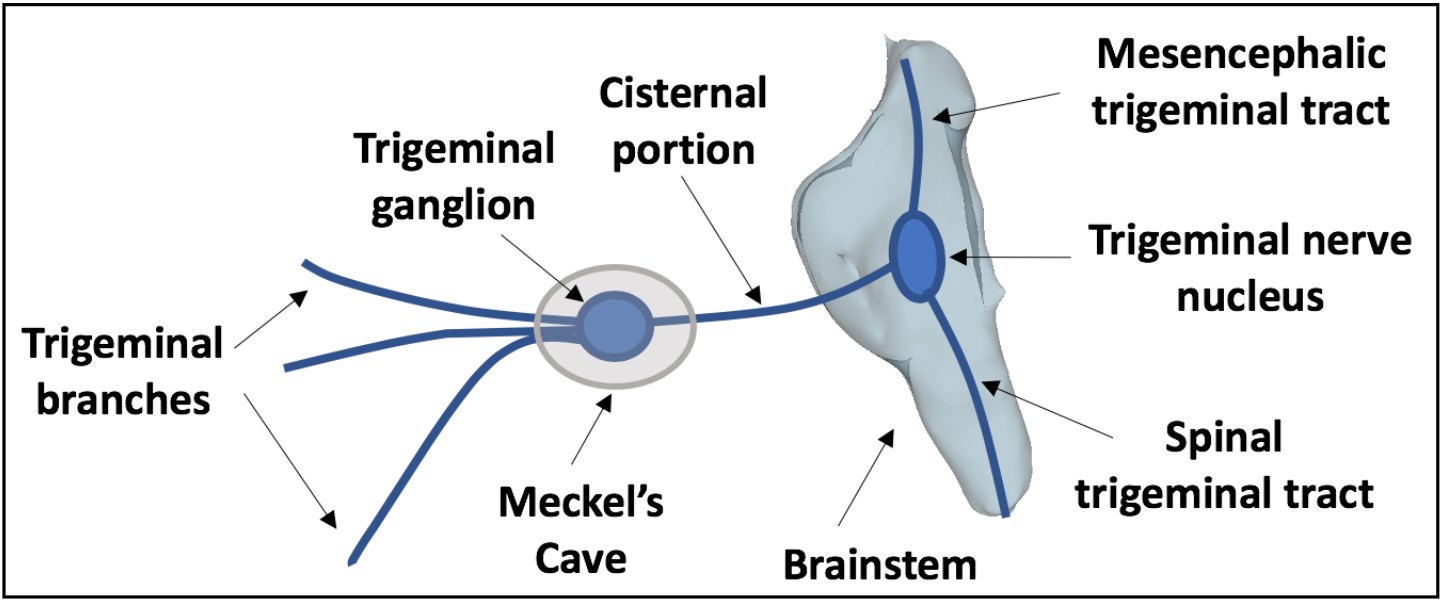
A schematic anatomical overview of the TGN.

Magnetic resonance imaging (MRI) techniques have been used to identify the TGN for clinical and research purposes (Casselman et al., 2008; Ciftci et al., 2004; Timothee Jacquesson et al., 2019; Ruiz-Juretschke et al., 2018; Tsutsumi et al., 2018; N. Yoshino et al., 2003). Among these techniques, traditional T2-weighted MRI is the most widely used, e.g., to confirm the presence of neurovascular compression at the root entry zone (REZ) of the TGN (Casselman et al., 2008; Xie et al., 2020). There have also been studies applying MRI techniques, such as constructive interference in steady-state sequence (CISS), fast imaging employing steady-state acquisition (FIESTA) and driven equilibrium radio frequency reset pulse (DRIVE), which have advanced performance in visualizing human nerves compared to a conventional T2-weighted image (Ciftci et al., 2004; Ruiz-Juretschke et al., 2018; Tsutsumi et al., 2018; N. Yoshino et al., 2003). However, these MRI sequences can only localize the cisternal portion of the TGN, while the continuity and pathological alteration of the TGN and brainstem nuclei, as well as the 3D relationship of the TGN with surrounding structures, cannot be assessed (Li et al., 2017; Liu et al., 2013; Neetu et al., 2016).

Diffusion MRI (dMRI), via a process called tractography, can track brain white matter and nerve fibers in vivo non-invasively based on the principle of detecting the random motion of water molecules in neural tissue (Basser et al., 1994, 2000). dMRI tractography has been applied successfully for tracking of the TGN (Fujiwara et al., 2011; Hung et al., 2017; Ishida et al., 2011; Timothée Jacquesson et al., 2018; Wei et al., 2016; M. Yoshino et al., 2016). One advantage of dMRI tractography is that it enables tracking of the 3D trajectory of the TGN for visualization of TGN structures not visualized by conventional MRI sequences (e.g., T2-weighted image, T2w), such as the course of the TGN within the brainstem as well as anterior to the cisternal portion (Timothée Jacquesson et al., 2018; Xie et al., 2020).

Currently, identification of the TGN from dMRI tractography data relies on the region of interest (ROI) selection strategy, where trained experts select TGNs in an interactive way by placing ROIs. In the literature, to our knowledge all related studies of the TGN have applied the expert ROI selection strategy (Behan et al., 2017; David Q. Chen, DeSouza, et al., 2016; David Qixiang Chen et al., 2011; Coskun et al., 2017; Fujiwara et al., 2011; Hung et al., 2017; Kabasawa et al., 2007; Moon et al., 2018; Wei et al., 2016; Xie et al., 2020; M. Yoshino et al., 2016; Zolal et al., 2017); however, practical problems remain. First, identification of the TGN is sensitive to ROI placement (Timothée Jacquesson et al., 2018; Xie et al., 2020), where selection of the best-performing ROIs is a challenge. In related work, ROI placement is variable across studies, where adopted ROIs include cisternal portion (CP, also called prepontine cistern, cisternal segment or midpoint of the cisternal segment), root entry zone (REZ), and/or the Meckel’s cave (MC) (Behan et al., 2017; David Q. Chen, DeSouza, et al., 2016; David Qixiang Chen et al., 2011; Coskun et al., 2017; Fujiwara et al., 2011; Kabasawa et al., 2007; Moon et al., 2018; Wei et al., 2016; Zolal et al., 2017). Second, placement of ROIs can be affected, or even fail, because of imaging artifacts and/or noise at the complex skull base environment (containing nerve, bone, air, soft tissue and cerebrospinal fluid) (Xie et al., 2020). Third, placement of ROIs may require inter-modality registration between dMRI and anatomical MRI (e.g. T2-weighted) data, which is challenging for dMRI with low image resolution (Malinsky et al., 2013) and echo-planar imaging (EPI) distortions (Albi et al., 2018). While ROI placement for TGN identification can be done using dMRI data directly (Behan et al., 2017; Fujiwara et al., 2011; Kabasawa et al., 2007; Moon et al., 2018; Xie et al., 2020), most studies have obtained ROIs from high-resolution anatomical MRI images for a better tissue contrast, requiring a co-registration to the low-resolution dMRI space (David Q. Chen, DeSouza, et al., 2016; David Qixiang Chen et al., 2011; Coskun et al., 2017; Fujiwara et al., 2011; Hung et al., 2017; Krishna et al., 2016; M. Yoshino et al., 2016; Zolal et al., 2017). Fourth, ROI placement depends critically on the experience of trained experts and hence inter-observer variability is a real and ongoing issue of accurate image interpretation (Hakulinen et al., 2012). Last but not least, manual interpretation is also time-consuming, inefficient and has clinical and expert labor costs.

In neuroscience, there has been an enduring interest in automated image processing and interpretation to resolve inter-observer variability and improve clinical efficiency, e.g., automatically locating brain anatomical structures and functions with references to common atlas spaces (Fischl, 2012; Maldjian et al., 2003). There are voxel-wise atlases that enable automated identification of cranial nerves in terms of the presence at a particular location in the brain (Fischl, 2012; Kikinis et al., 1996; Sultana, 2017). However, these atlases cannot be used to automatically identify tractography fibers belonging to the cranial nerves. Another type of voxel-wise atlas can define ROIs that are useful for selecting tractography fibers. While this approach has mainly been applied in the cerebrum (Lawes et al., 2008; Y. Zhang et al., 2010), Chen et al. demonstrated successful automated subject-specific identification of several cranial nerves (the facial/vestibular-cochlear nerve complex and the vagus nerve) using a voxel-wise ROI atlas (David Q. Chen, Zhong, et al., 2016). However, such ROI-based methods can be challenged by highly sensitive tractography methods, which require more ROIs to select due to their increased sensitivity (O’Donnell et al., 2017; Xie et al., 2020). Rather than creating a voxel-wise atlas, in brain white matter analysis, many studies have created brain dMRI tractography atlases (Guevara et al., 2017; Maddah et al., 2005; O’Donnell & Westin, 2007; Román et al., 2017; Yoo et al., 2015; Fan Zhang, Wu, et al., 2018; Ziyan et al., 2009). These studies have successfully demonstrated automated identification of anatomical white matter fiber tracts (e.g. arcuate fasciculus), with several advantages including 1) consistent tract identification in the dMRI data from different acquisition protocols, 2) using dMRI data only, thus not requiring inter-modality registration, and 3) high efficiency to reduce expert labor costs and enable tractography analysis in large-scale dMRI datasets. However, to our knowledge there are no existing tractography atlases that can enable automated identification of the TGN in tractography.

In this study, we present what we believe is the first study to create a dMRI tractography TGN atlas, which enables automated identification of the TGN in new tractography data without relying on expert ROI placement. Our method relies on a well-established groupwise fiber clustering pipeline from our research group (O’Donnell et al., 2012; O’Donnell & Westin, 2007), which has been successfully applied in multiple research studies (Fan et al., 2019; Gong et al., 2018; O’Donnell et al., 2017; Stojanovski et al., 2019; Wu et al., 2018; Fan Zhang, Savadjiev, et al., 2018; Fan Zhang, Wu, et al., 2018) and has been used recently for creation of an anatomically curated white matter tract atlas (Fan Zhang, Wu, et al., 2018, 2019). In the present study, we employ this fiber clustering pipeline to identify common TGN structures in an atlas population, including 50 subjects from the Human Connectome Project (HCP) (Van Essen et al., 2013) that provide high-quality dMRI data. Leveraging population-based brain anatomical information and expert neuroanatomical knowledge, we identify a total of 40 fiber clusters belonging to the TGN in the atlas. Each cluster represents a certain anatomical subdivision of the TGN and its variability in the atlas population. The curated TGN model includes not only the cisternal portion but also the putative mesencephalic tract (Shigenaga et al., 1989) and the putative spinal trigeminal tract (M. Yoshino et al., 2016), which are important portions of the TGN but have been relatively less studied. The created TGN atlas and the fiber clustering pipeline also provide a method to automatically identify the TGN in new subject datasets. We demonstrate a successful application to dMRI datasets from two different acquisition sites, including those from a clinical acquisition protocol (a lower spatial resolution than the HCP data). The fiber clustering pipeline is open source^1^ and the TGN atlas will be made available online^2^, as part of the SlicerDMRI project^3^ (Norton et al., 2017; Fan Zhang et al., 2020).

In the rest of the paper, we first describe the datasets in this study. Here, we leveraged several findings about data processing, TGN tracking and ground truth identification from our previous work, where we compared different TGN fiber tracking strategies (dMRI data with different b-values, in combination with both single- and multi-tensor tractography methods) (Xie et al. 2020). Then, we introduce the creation of the proposed TGN atlas from 50 healthy adults, followed by a demonstration of our method with an application of the atlas to dMRI datasets that were scanned at two different acquisition sites (with different spatial and angular resolutions). Quantitative and qualitative evaluations are performed to evaluate our method’s TGN identification performance, with comparison to expert selected TGNs.

## 2. Methods

### 2.1 Datasets and data processing

#### 2.1.1 Datasets

In this study, we used dMRI data from two acquisition sites, including the HCP database (Van Essen et al., 2013) and the Parkinson’s Progression Markers Initiative (PPMI) (Marek et al., 2011) database. The HCP data was used for the TGN atlas creation (50 atlas subjects) and experimental evaluation (an independent set of 50 testing subjects), while the PPMI data (40 subjects) was used only for experimental evaluation. Table 1 gives an overview of the demographics and the dMRI acquisitions of the HCP and PPMI datasets under study.

**Table 1:**
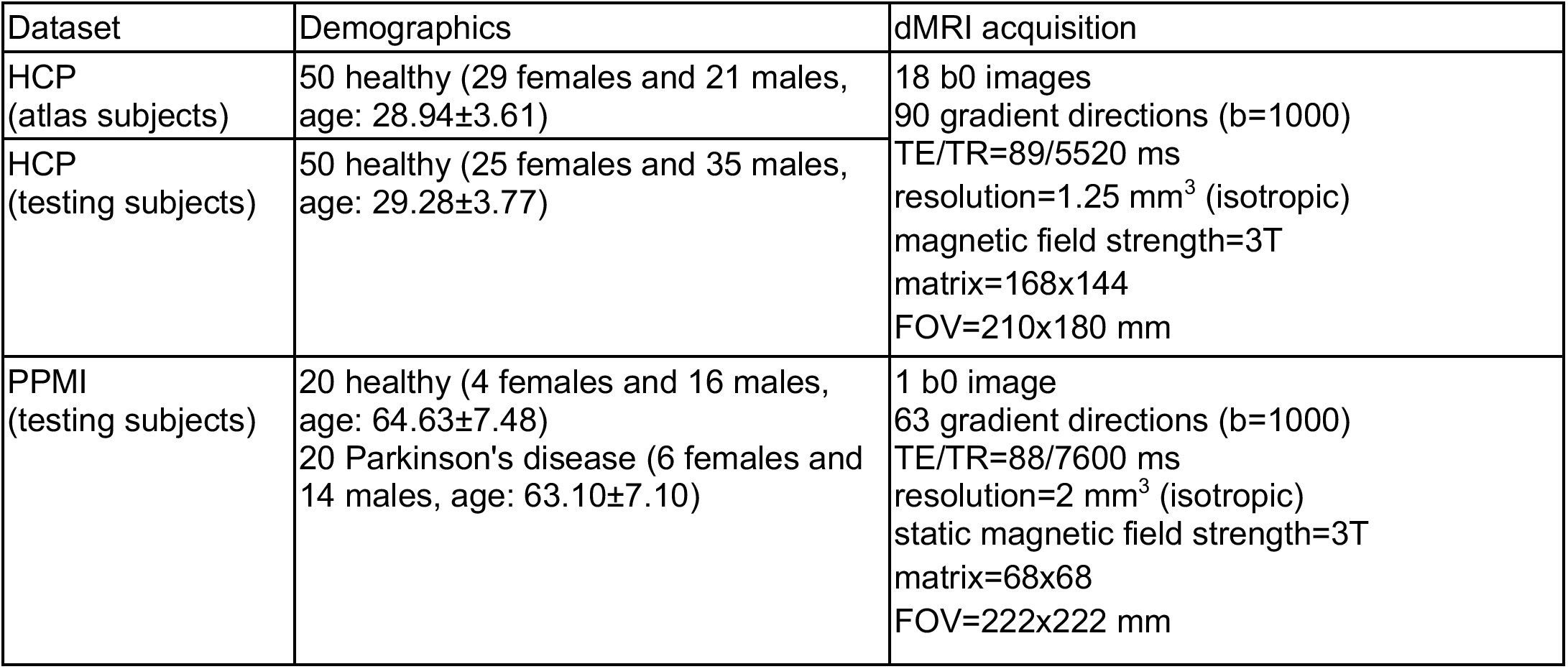
Demographics and dMRI acquisition of the HCP and PPMI datasets under study.

The HCP provides dMRI data that was acquired with a high quality image acquisition protocol using a customized Connectome Siemens Skyra scanner and processed using a well-designed processing pipeline (Glasser et al., 2013) including motion correction, eddy current correction and EPI distortion correction. The acquisition parameters of the dMRI data in HCP were: TE=89.5 ms, TR=5520 ms, and voxel size=1.25×1.25×1.25 mm^3^. A total of 288 images were acquired in each dMRI dataset, including 18 baseline images with a low diffusion weighting b=5 s/mm^2^ and 270 diffusion-weighted (DW) images evenly distributed at three shells of b=1000/2000/3000 s/mm^2^. More detailed information about the HCP data acquisition and preprocessing can be found in (Glasser et al., 2013). In our study, we used the single-shell b=1000 s/mm^2^ data to perform TGN tracking (see Section 2.1.2 for details) because it represents the clinical acquisition protocol and was shown in our previous study to be more effective for TGN identification than higher b values (Xie et al., 2020). We also used the anatomical T2w data for evaluation of the TGNs. The acquisition parameters used for the T2w data were TE=565 ms, TR=3200 ms, and voxel size=0.7×0.7×0.7 mm^3^. Imaging data from a total of 100 HCP subjects was used in our study, including 50 subjects for the TGN atlas creation and another 50 subjects for experimental evaluation. We note that to ensure high quality TGN representations for atlas creation, we selected 50 atlas subjects whose dMRI data did not have apparent imaging artifacts and/or noise. (In our previous work (Xie et al., 2020), we showed that manual TGN identification failed in several of the 100 HCP subjects because of imaging artifacts and/or noise at the skull base region.)

The PPMI data was used to test TGN identification performance using an acquisition protocol that was different from the HCP data. We chose data from Parkinson’s disease because it has been suggested to be closely related to the TGN (Tremblay et al., 2017). The acquisition parameters of the dMR data were: TE=88 ms, TR=7600 ms, voxel size=2×2×2 mm^3^, 1 baseline image with b=0 s/mm^2^ and 64 DW images with b=1000 s/mm^2^. T2-weighted data (co-registered with the dMR data) was also used for TGN experimental evaluation. The acquisition parameters for the T2w data were: TE=101 ms, TR=3000 ms, and voxel size=1×1×1 mm^3^. The dMRI data was pre-processed with the following steps. Eddy current-induced distortion correction and motion correction were conducted using the Functional Magnetic Resonance Imaging of the Brain (FMRIB) Software Library tool (Jenkinson et al., 2012). An echo-planar imaging (EPI) distortion correction was performed with reference to the T2-weighted image using the Advanced Normalization Tools (ANTS) (Avants et al., 2009). For each subject, a nonlinear registration (registration was restricted to the phase encoding direction) was computed from the b0 image to the T2w image to make an EPI corrective warp. Then, the warp was applied to each diffusion image. Data from 40 PPMI subjects (20 healthy controls and 20 Parkinson’s disease patients) was used in our experiment.

#### 2.1.2 Multi-tensor TGN tractography

For each subject under study, we performed TGN tractography from the dMRI data. We used the two-tensor unscented Kalman filter (UKF) tractography method^4^ (Malcolm et al., 2010; Reddy & Rathi, 2016) to perform TGN tracking, as illustrated in Figure 2(a). We chose the two-tensor UKF tractography method because it has been demonstrated to be effective in tracking the TGN in our previous study (Xie et al., 2020), as well as tracking the brain white matter fiber tracts (Z. Chen et al., 2016; Gong et al., 2018; Liao et al., 2017; Fan Zhang, Wu, et al., 2018). The UKF method fits a mixture model of two tensors to the dMRI data while tracking fibers, providing a highly sensitive fiber tracking ability, in particular, in the presence of crossing fibers (Z. Chen et al., 2016; Gong et al., 2018; Liao et al., 2017; Fan Zhang, Wu, et al., 2018). This is important for tracking the intra-brainstem portions of the TGN (including the putative spinal trigeminal and putative mesencephalic trigeminal tracts, as demonstrated in our previous study (Xie et al., 2020) and in Supplementary Figure 2), where nerve fibers cross with white matter fibers. In contrast to other methods that fit a model to the signal independently at each voxel (Behan et al., 2017; Qazi et al., 2009), the UKF method fits a model to the diffusion data while tracking fibers, in a recursive estimation fashion (the current tracking estimate is guided by the previous one). One benefit of the recursive estimation is to help stabilize model fitting; thus fiber tracking can be robust to a certain amount of imaging artifact/noise. Another benefit of UKF is that fiber tracking orientation is controlled by a probabilistic prior about the rate of change of fiber orientation (defined as the parameter Qm introduced below), instead of cutoffs or limits on the fiber curvature as in typical tractography algorithms. Consequently, sharp fiber curvatures are avoided as they are very unlikely, whereas fiber curvatures (e.g., the branching structures of the TGN) supported by the dMRI are still allowed. These properties of UKF are different from other tractography algorithms such as the single DTI tractography (Basser et al., 2000) and the two-tensor eXtended Streamline Tractography (Qazi et al., 2009) that have been applied for TGN fiber tracking (David Q. Chen, DeSouza, et al., 2016; David Qixiang Chen et al., 2011; Coskun et al., 2017; Kabasawa et al., 2007). (To demonstrate our automated TGN identification method’s ability to generalize to tractography data from different methods, we have included an additional two tractography methods, as introduced in Supplementary Material S1.)

**Figure 2.**
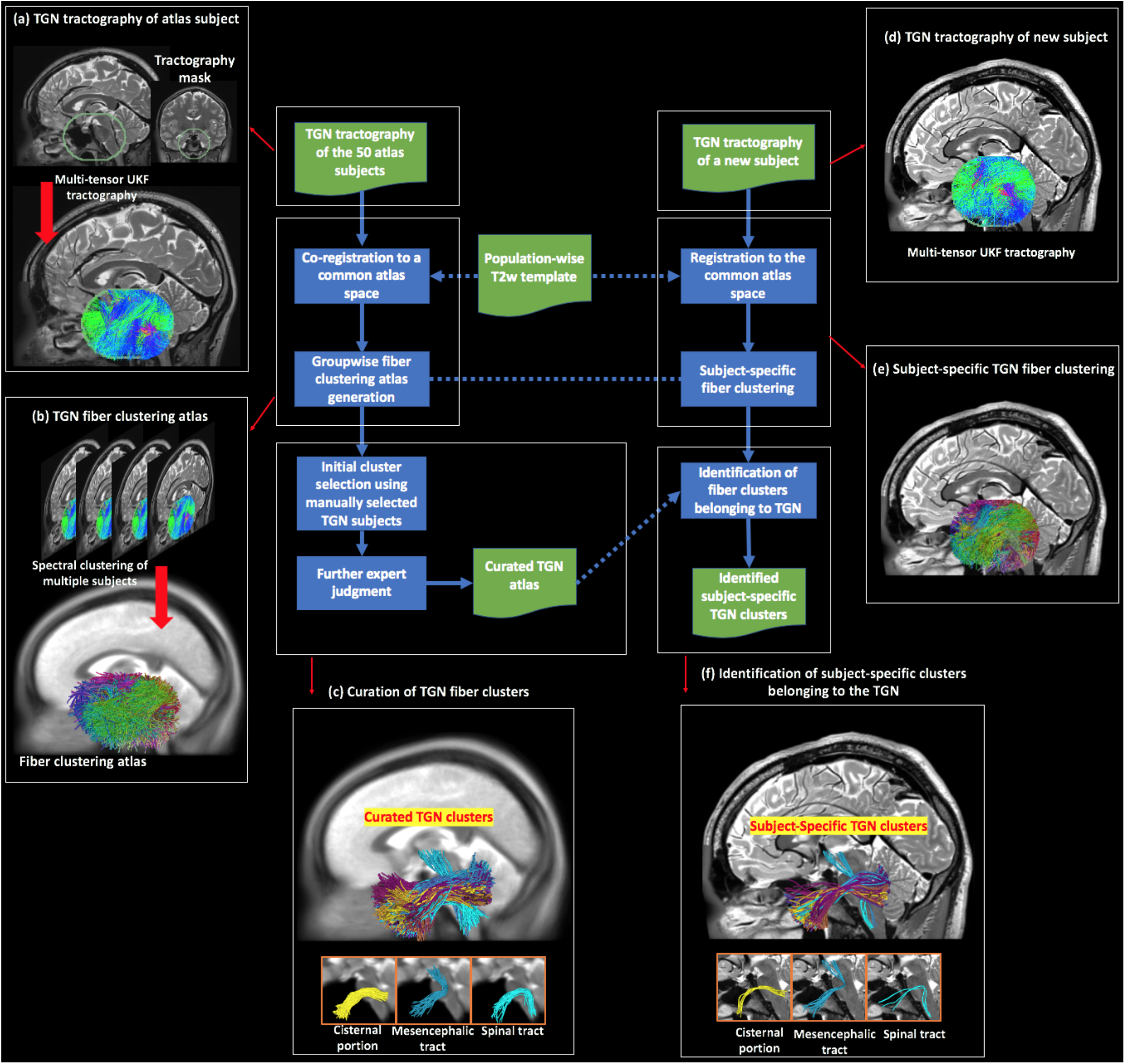
Schematic diagram of the proposed method, where the blue boxes represent the computational steps and the green boxes represent the input/output. The major steps are linked to a pictorial illustration, as follows. (a) to (c) show the creation of the TGN atlas. (a) Multi-tensor tractography is seeded within a mask that covers the possible region through which the TGN passes. (b) Given the tractography data (co-registered to a common space, i.e., the atlas space) from the 50 atlas subjects, spectral clustering is performed to generate a fiber clustering atlas, where each cluster has a unique color. (c) Using expert neuroanatomical knowledge (involving three experts, GX, MAM, and NM), 40 clusters were identified to belong to the TGN, where each cluster represents a specific subdivision of the whole TGN. Three example clusters are displayed, belonging to the cisternal portion, the mesencephalic trigeminal tract, and the spinal trigeminal tract, respectively. (d) to (f) show identification of the TGN of a new subject. (d) TGN tractography of the new subject is performed, in the same way as the atlas subjects. (e) Fiber clustering of the tractography data (registered to the atlas space) is conducted according to the fiber clustering atlas. (f) Identification of the TGN clusters in the new subject is conducted by finding the corresponding subject-specific clusters to those annotated in the atlas. Three example subject-specific TGN clusters, corresponding to the ones shown in (c), are displayed.

TGN tractography was seeded from all voxels within a mask, which was larger than the possible region through which the TGN passes. This procedure was similar to whole brain seeding but it was restricted to the potential TGN region for efficiency. We note that our method does not require sophisticated masking as long as the mask covers the TGN and is approximately in a similar place across all subjects. We used the 3D Slicer Segment Editor tool to do this by placing a spherical or oval mask with a diameter about 35mm, centered at the anterior portion of the pons (as illustrated in Figure 2(a)). There are five major parameters of the UKF method, including *seedingFA*, *stoppingFA*, *stoppingThreshold, Qm* and *Ql.* These parameters function as follows. Tractography is seeded in all voxels within a provided mask where fractional anisotropy (FA) is greater than *seedingFA.* Tracking stops in voxels where the FA value falls below *stoppingFA* or the sum of the normalized signal across all gradient directions falls below *stoppingThreshold* (a parameter to distinguish between white/gray matter and cerebrospinal fluid (CSF) regions). During the tracking, the UKF method uses *Qm* to control process noise for angles/direction, and *Ql* to control process noise for eigenvalues. These UKF tractography parameters were well tuned and were set as: *seedingFA* = 0.06, *stoppingFA* = 0.05, *stoppingThreshold* = 0.06, *Qm* = 0.001 and *Ql* = 300. Two seeds per voxel were used for seeding the tractography, which resulted in about 70,000 fibers in the TGN tractography per subject. Visual and quantitative quality control of the tractography data was performed using the quality control tool in the *whitematteranalysis*^5^ software. We note that in the present study, we used a relatively low seeding sampling setting (2 seeds per voxel) because it was sufficient to generate visually reasonable TGNs corresponding to the anatomy, while keeping a low computational cost. To increase the tract density, a higher number of seeds per voxel can be used (see Supplementary Material S1 for a visualization of an identified TGN from tractography data computed using 5 seeds per voxel).

#### 2.1.3. Identification of ground truth TGN using manual selection

For selected subjects, we performed manual ROI-based TGN identification from the tractography data. These manually selected TGNs were used for initial selection of fiber clusters potentially belonging to the TGN in the atlas (see Section 2.2.1) and were used as ground truth for evaluation of the automatically identified TGNs (see Section 2.4). (We note that for identification of the TGN in new subjects using our method, there is no need to perform manual TGN selection.)

We performed manual TGN identification using predefined manually drawn ROIs from the MC and the CP of the TGN, as described in (Xie et al., 2020). We note that these two ROIs were most commonly used in the literature for expert TGN selection (Behan et al., 2017; Coskun et al., 2017; Fujiwara et al., 2011; Kabasawa et al., 2007; Wei et al., 2016; Xie et al., 2020; M. Yoshino et al., 2016; Zolal et al., 2017), and they were relatively easily identified. The manual TGN identification method was previously validated in an inter-rater experiment (by two practicing neurosurgeons GX and MAM), showing a high joint probability of agreement (Xie et al., 2020). The ROI in MC was drawn on the mean b=0 image from the coronal view, and the ROI in CP was drawn on the mean directionally encoded color (DEC) map of diffusion tensor imaging (DTI) from the coronal view. For the HCP data, we attempted to perform manual TGN selection on all of the 100 subjects; however, 8 testing subjects failed because their dMRI data had artifacts and/or noise at the skull base region that prevented placement of ROIs. (We note that we also attempted to draw ROIs on the T2w image, on which the ROIs could be recognized. However, this attempt also failed because the imaging artifacts and/or noise affected the registration between the dMRI and T2w data at the skull base region.) For the PPMI database, we performed manual TGN identification on two randomly selected subjects (a 69 year old female healthy control and a 72 year old male Parkinson’s disease patient).

### 2.2. Creation of TGN atlas

We created the TGN fiber clustering atlas using the tractography data from the 50 atlas subjects. This involved *1)* generating a data-driven fiber clustering atlas for TGN tractography parcellation into multiple fiber clusters and *2)* curating fiber clusters anatomically belonging to the TGN, as illustrated in Figure 2(b,c).

#### 2.2.1 Generation of TGN fiber clustering atlas

The TGN fiber clustering atlas was generated using groupwise fiber clustering to simultaneously parcellate TGN tractography from multiple subjects (Figure 2(b)). First, the TGN tractography data of the 50 atlas subjects was registered into a common space (i.e. atlas space). This was done by performing an affine registration between the b=0 image of each subject (moving image) and a population-mean T2-weighted image (reference image) using 3D Slicer. We chose T2w data for co-registration because it has similar contrast to the b=0 images for promising inter-MRI-modality registration (Albi et al., 2018). Also, T2w data has a good contrast of the cisternal portion of the TGN and has been widely used to confirm the presence of the TGN (Casselman et al., 2008; Xie et al., 2020). Specifically, we used the population-mean T2 image, that has been successfully applied to co-register tractography data (Fan Zhang, Hoffmann, et al., 2019), provided in the white matter atlas from our group (Fan Zhang, Wu, et al., 2018). Then, the obtained transform was applied to the subject-specific TGN tractography data. In the present study, we performed a semi-automated quality control of the registration results, using in-house developed Matlab scripts that enable a visualization of the registered b0 and the population-mean T2 image together.

Next, spectral clustering was used to compute a high-dimensional fiber clustering atlas (O’Donnell & Westin, 2007) to divide the TGN tractography into *K* clusters, where *K* is a user-given parameter to define the parcellation scale. The spectral embedding created a space that robustly represented each fiber according to its affinity to all other fibers across subjects. This fiber representation gives a robust feature vector or "fingerprint" that describes the fiber for clustering. The fiber affinity was computed by converting pairwise fiber geometric distances (the popular mean closest point distance is used (Moberts et al., 2005; O’Donnell & Westin, 2007)) using a Gaussian-like kernel, representing fiber similarity according to the fiber geometry and trajectory. One benefit of such fiber similarity matching was that it was highly robust to local fiber tract variation to ensure morphology agreement across subjects. Therefore, roughly aligned tractography data using the above volume-based affine co-registration was sufficient to co-register across different subjects. Nystrom sampling (Fowlkes et al., 2004) was used to reduce the computations considering the large number of fiber pairs across subjects. Bilateral clustering, simultaneously segmenting TGN fibers on both sides of the cranial base to improve parcellation robustness (O’Donnell & Westin, 2007), was applied to obtain the *K* fiber clusters. Bilateral clustering is beneficial for accounting for potential laterality of the TGNs because it can robustly find the corresponding fiber structures on both sides of the cranial base. Our previous studies have demonstrated the benefit of the bilateral fiber clustering in identifying corresponding white matter structures across hemispheres and in investigating potential white matter lateralized changes (Propper et al., 2010; Wu et al., 2018). In addition, we incorporated an outlier removal process to remove improbable fibers for cluster consistency in the atlas. In this process, a fiber was considered as an outlier if it was distant from other fibers within its cluster (over 2 standard deviations from the cluster’s mean fiber affinity, as applied in our previous work (O’Donnell et al., 2017; Wu et al., 2018; Fan Zhang, Savadjiev, et al., 2018; Fan Zhang, Wu, et al., 2018)). In the present study, all fiber clustering computations were performed using the *whitematteranalysis* software, with the suggested settings for related parameters. 10,000 fibers were randomly sampled from each subject’s TGN tractography for a total of 0.5 million fibers for the atlas creation.

We generated multiple fiber clustering atlases to investigate the TGN tractography parcellation at different scales (number of clusters, *K*, ranging from 500 to 3000). The TGN atlas consisting of *K*=2500 clusters was chosen because it represented the minimum scale to identify the putative mesencephalic trigeminal tract. The putative mesencephalic trigeminal tract was a small TGN structure with fewer fibers than the TGN cisternal portion. Using a coarser parcellation scale (e.g. *K*=2000), the putative mesencephalic trigeminal tract was clustered together with other TGN structures. On the other hand, while using a finer parcellation scale (e.g. *K*=3000) could also provide a reasonable TGN clustering result, it would increase the workload for expert curation of the TGN clusters and decrease parcellation consistency (i.e. consistent identification of each individual cluster) across subjects as suggested in our previous work (F. Zhang, Norton, et al., 2017; Fan Zhang, Wu, et al., 2018).

#### 2.2.2 Curation of TGN fiber clusters

Given the chosen TGN fiber clustering atlas (*K*=2500), each fiber cluster (including bilateral fibers on both sides of the cranial base) was annotated to indicate whether it belongs to the TGN or not. This fiber cluster annotation was performed by an initial cluster annotation computation, followed by expert judgment. This resulted in a curated atlas TGN with multiple fiber clusters, where nerve fibers in each cluster have similar trajectories, representing a particular anatomical subdivision. In the way, the curated TGN atlas provides an effective way to describe the complex anatomy of the TGN, e.g., the branching structures of the TGN that are subdivided into multiple clusters.

We leveraged the manually selected TGNs (Section 2.1.3) to perform an initial selection of potential clusters belonging to the TGN. The purpose of this initial computation step is to bootstrap the expert cluster annotation with a first pass that can be performed automatically by the computer. From the 10,000 fibers that were randomly sampled from each subject’s TGN tractography for the atlas generation, we first identified the fibers that were manually selected to be the TGNs. Then we calculated a probability for each bilateral atlas cluster belonging to the TGN, i.e., the number of fibers that were manually selected to belong to the TGN divided by the number of total fibers in the cluster. We initially selected the fiber clusters that had a probability over 0, which resulted in a total of 127 candidate clusters for expert judgment.

Next, an expert rater (GX who is a practicing neurosurgeon) performed expert annotation of the 127 candidate bilateral clusters. Specifically, the expert rater viewed each atlas fiber cluster with reference to the population-mean T2 image that was used to register all atlas TGN tractography into the atlas space (Section 2.2). This enabled viewing of the nerve structure and its variability across all atlas subjects. To confirm the population-based decision, the corresponding subject-specific clusters from five randomly selected subjects were checked with reference to the subject’s T2-weighted image. Another expert rater (NM who is a neuroanatomist) viewed the curated TGN clusters and confirmed their anatomical correctness.

Overall, there are a total of 40 TGN clusters in the atlas (Figure 2(c)). Each cluster represents a particular anatomical subdivision of the TGN, including the TGN cisternal portion (35 clusters), the putative mesencephalic tract (2 clusters), and the putative spinal trigeminal tract (3 clusters) (example clusters are shown in Figure 2(c)).

### 2.3 Application of the TGN atlas to new subjects

Automated identification of the TGN of a new subject was conducted by applying the atlas to the subject’s TGN tractography, as illustrated in Figure 2(d,e,f). First, the TGN tractography was registered into the atlas space, by performing an affine registration between the subject’s b=0 image and the population-mean T2-weighted image and then applying the obtained transform to the TGN tractography data. (This was the same process as registering the TGN tractography data of the atlas subjects.) Then, subject-specific fiber clusters were detected using spectral embedding of the registered tractography, followed by assignment of each fiber to the closest atlas cluster (O’Donnell & Westin, 2007). As a result, the new subject’s TGN tractography was divided into multiple fiber clusters, where each cluster corresponded to a certain atlas fiber cluster. Outlier fibers were removed if their fiber affinity regarding the atlas cluster was over 2 standard deviations from the cluster’s mean fiber affinity. Next, TGN identification of the new subject was conducted by automatically finding the subject-specific clusters that corresponded to the annotated atlas clusters.

### 2.4 Experimental evaluation

All subjects’ TGN tractography (including the 50 HCP atlas subjects, the 50 HCP testing subjects and the 40 PPMI subjects) was parcellated using the proposed atlas. We note that our atlas was created using only the 50 HCP atlas subjects, while the additional 50 HCP testing subjects and the 40 PPMI subjects were used for testing the performance on new data. We performed the following experiments to evaluate the TGN identification performance of our method.

#### 2.4.1 TGN identification rate

We computed the mean TGN identification rate (percentage of successfully identified TGNs) across all subjects in each dataset. We performed this evaluation for the overall TGN, as well as for the subdivisions including the cisternal portion of the TGN, the mesencephalic trigeminal tract and the spinal trigeminal tract. The identified TGNs and their subdivisions were confirmed (i.e., visually assessed as belonging to the TGN) by the expert rater (GX). We note that this expert visual inspection provided a complementary assessment to ground truth comparison with manual ROI selection, which is especially important to ensure the anatomical viability of the automatically identified TGNs when manual ROI selection failed. We reported the mean identification rate of the 50 HCP testing subjects (a total of 100 TGNs) and the 40 PPMI testing subjects (a total of 80 TGNs). For comparison, we also reported the mean identification rate of the 50 HCP atlas subjects (a total of 100 TGNs) to show how the atlas generalized to data from the atlas population. Finally, we reported the mean TGN identification rates of the expert TGN selection in the 50 HCP atlas subjects and in the 50 HCP testing subjects.

#### 2.4.2 TGN spatial overlap

We performed a quantitative comparison to assess if the TGNs identified using the atlas spatially overlapped with the manually identified TGNs. Specifically, we computed the weighted Dice (wDice) coefficient between the automatically and manually identified TGNs from each subject. wDice coefficient was designed specifically for measuring volumetric overlap of fiber tracts (Cousineau et al., 2017; Fan Zhang, Wu, et al., 2019). wDice extends the standard Dice coefficient (Dice, 1945) taking account of the number of fibers per voxel so that it gives higher weighting to voxels with dense fibers. For the HCP database, we reported the mean and the standard deviation of the wDice values across the 50 atlas subjects and those across the 42 testing subjects with successful manual TGN selection. (Note that the other 8 HCP testing subjects were not included in the quantitative comparison because manual selection failed.) For the PPMI database, we reported the wDice score for each of the two selected subjects with manually selected TGNs.

#### 2.4.3 TGN visualization

For visual comparison of the automated TGN identification and the manual selection, we rendered the automatically and manually identified TGNs from three example subjects. These included one HCP testing subject with successful manual TGN identification, one HCP testing subject with unsuccessful manual TGN identification, and one PPMI testing subject (healthy control). We also provided a visualization to demonstrate the effects of the dMRI imaging artifacts and/or noise on the ROI placement in the HCP testing subject with unsuccessful manual TGN identification.

We then provided a visualization of TGNs to show the anatomical regions where the TGN passed. We first showed the curated TGN in the atlas. This was done by rendering a voxel-based fiber density heatmap that quantifies the number of fibers present in each voxel and the regions through which the TGN passed on the population-mean T2-weighted image (used for co-registering the TGN tractography data). We also showed subject-specific TGNs, by rendering their fiber density heatmaps and the regions through which the TGN passed on the corresponding subject T2w images. Three example subjects (the same subjects as used in the above visual comparison) were used in this visualization.

## 3. Results

### 3.1 TGN identification rate

All TGNs were successfully identified using the proposed automated identification method in all subjects under study, including the 50 HCP atlas subjects, the 50 HCP testing subjects (including the 8 subjects where manual selection failed), and the 40 PPMI testing subjects (Table 2). Regarding the different subdivisions, the cisternal portion was identified in all subjects under study. We also obtained relatively high identification rates for the mesencephalic trigeminal tract (on average 50.0%) and spinal trigeminal tract (on average 52.4%) across all 100 HCP subjects, where the manual selection could successfully identify 34.5% of the mesencephalic trigeminal tracts and 37.0% of the spinal trigeminal tracts. For the PPMI data, the identification rates of the mesencephalic trigeminal and spinal trigeminal tracts using the proposed automated method were relatively low compared to those in the HCP data.

**Table 2.**
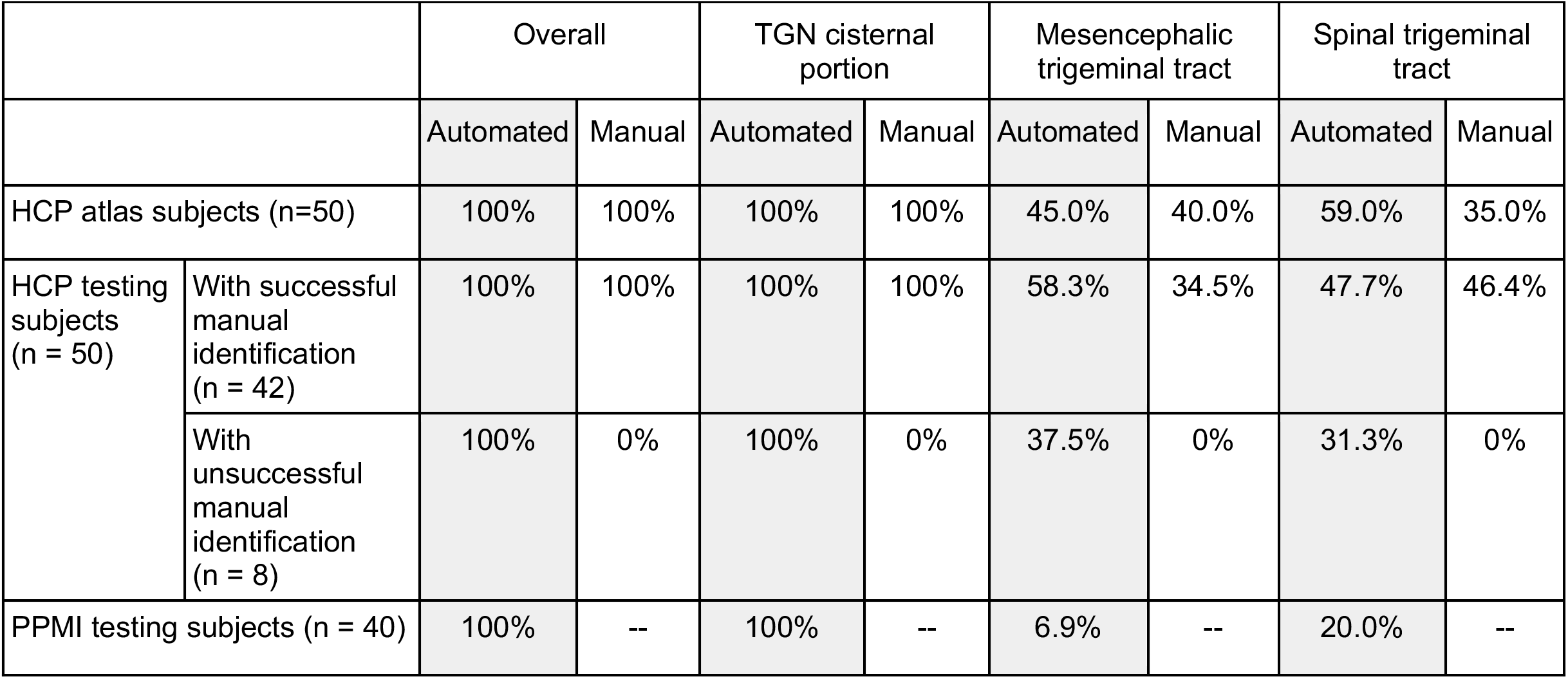
TGN identification rate (percentage of successfully identified TGNs) of the overall TGN and its subdivisions using the proposed automated identification method (highlighted in gray) and the manual selection method. For the PPMI data, we did not perform manual TGN identification; thus, the TGN identification rate was not reported.

|  |  | Overall |  | TGN cisternal portion |  | Mesencephalic trigeminal tract |  | Spinal trigeminal tract |  |
| --- | --- | --- | --- | --- | --- | --- | --- | --- | --- |
|  |  | Automated | Manual | Automated | Manual | Automated | Manual | Automated | Manual |
| HCP atlas subjects (n=50) |  | 100% | 100% | 100% | 100% | 45.0% | 40.0% | 59.0% | 35.0% |
| HCP testing subjects (n = 50) | With successful manual identification (n = 42) | 100% | 100% | 100% | 100% | 58.3% | 34.5% | 47.7% | 46.4% |
|  | With unsuccessful manual identification (n = 8) | 100% | 0% | 100% | 0% | 37.5% | 0% | 31.3% | 0% |
| PPMI testing subjects (n = 40) |  | 100% | -- | 100% | -- | 6.9% | -- | 20.0% | -- |

### 3.2 TGN spatial overlap

Table 3 gives the mean and the standard deviation of the wDice scores across the 50 HCP atlas subjects and those of the 42 HCP testing subjects with successful manual TGN identification. High mean wDice scores, on average over 0.75, were obtained. The threshold for a good wDice score to evaluate tract spatial overlap was suggested to be 0.72 according to Cousineau et al (Cousineau et al., 2017). Table 3 also gives the wDice scores of the two PPMI testing subjects with manually selected TGNs, which were about 0.79 for both subjects.

**Table 3.** Spatial overlap (wDice score) between automatically (proposed) and manually identified TGNs.

|  |  |  |
| --- | --- | --- |
| 50 HCP atlas subjects |  | 0.7585 ± 0.0883 |
| 42 HCP testing subjects (with successful manual identification) |  | 0.7534 ± 0.1127 |
| PPMI testing subjects | Subject 1 | 0.7892 |
|  | Subject 2 | 0.7887 |

### 3.3 TGN visualization

Figure 3 gives a visual comparison between the automatically (proposed) and manually identified TGNs for three example subjects. Highly visually comparable results were obtained between the two methods for the two subjects with successful manual identification. Our proposed method could also successfully identify a visually reasonable TGN on the subject where manual identification failed. In the testing subject with unsuccessful manual TGN identification, the skull base region is affected by imaging imaging artifacts and/or noise, where the predefined ROIs at the Meckel’s Cave and the cisternal portion were not visually accessible. (To confirm the anatomical validity of the automatically identified TGN in the testing subject, we have performed another manual TGN identification method that is based on interactively moving ROIs. A visualization of the TGNs is provided in Supplementary Figure 3.)

**Figure 3.**
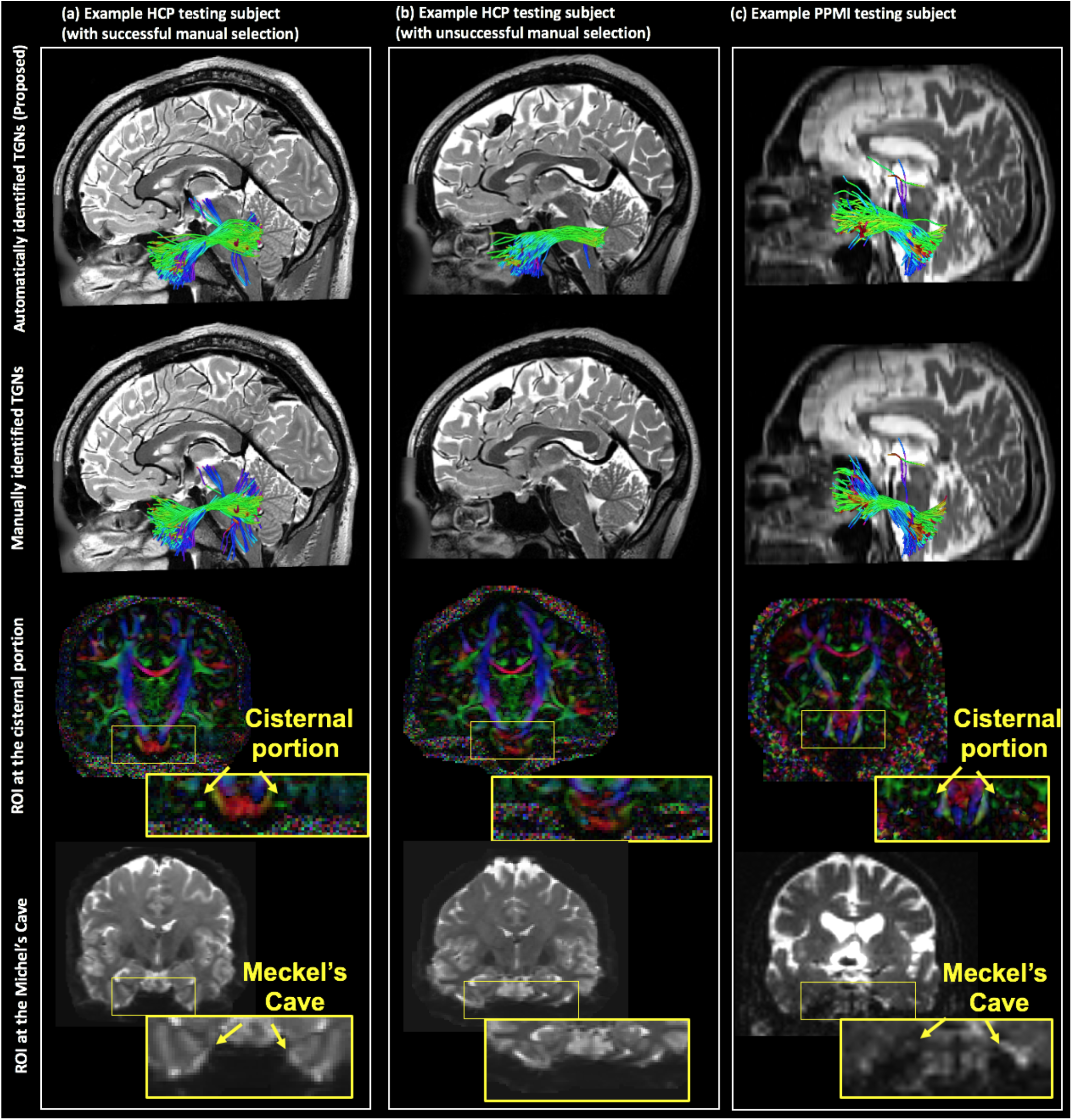
Visual comparison of the TGN 3D fiber trajectory between the automatically (proposed) and manually identified TGNs. The three example subjects include one HCP testing subject with successful manual TGN identification, one HCP testing subject with unsuccessful manual TGN identification, and one PPMI testing subject (healthy control). For the HCP testing subject with unsuccessful manual TGN identification, the b0 and DTI image are distorted, preventing successful placement of the ROIs in CP and MC.

Figure 4(a) shows the 3D fiber trajectory and the fiber density map of the TGN curated in the atlas, overlaid on the population mean T2w image. In general, the TGN had an anatomically correct shape and corresponded well to the known anatomy of the TGN pathways, i.e., passing through the Meckel’s Cave (MC) and overlapping well with the cisternal portion (CP), as appearing on the T2w image. Figure 4(b) gives the TGN visualization of the example HCP subject with successful manual selection. The TGNs identified using our method were anatomically correct, passing through the MC and overlapping with the CP as appearing on the T2w image. Figure 4(c) renders the identified TGNs from the example HCP subject with unsuccessful manual selection. In this subject, our obtained TGNs are visually anatomically correct in terms of the shape; however, they do not pass through the MC and do not overlap with the CP as appearing on the T2w data. This was because the dMRI data of this subject had imaging artifacts in the skull base region, which affected the registration with the T2w data at the skull base region. (This also explained the failure of our attempt to perform manual selection using ROIs drawn on the T2w data.) Figure 4(d) displays the TGNs identified from the example PPMI subject. The identified TGNs were anatomically correct, passing through the MC and overlapping with the CP as appearing on the T2w data.

**Figure 4.**
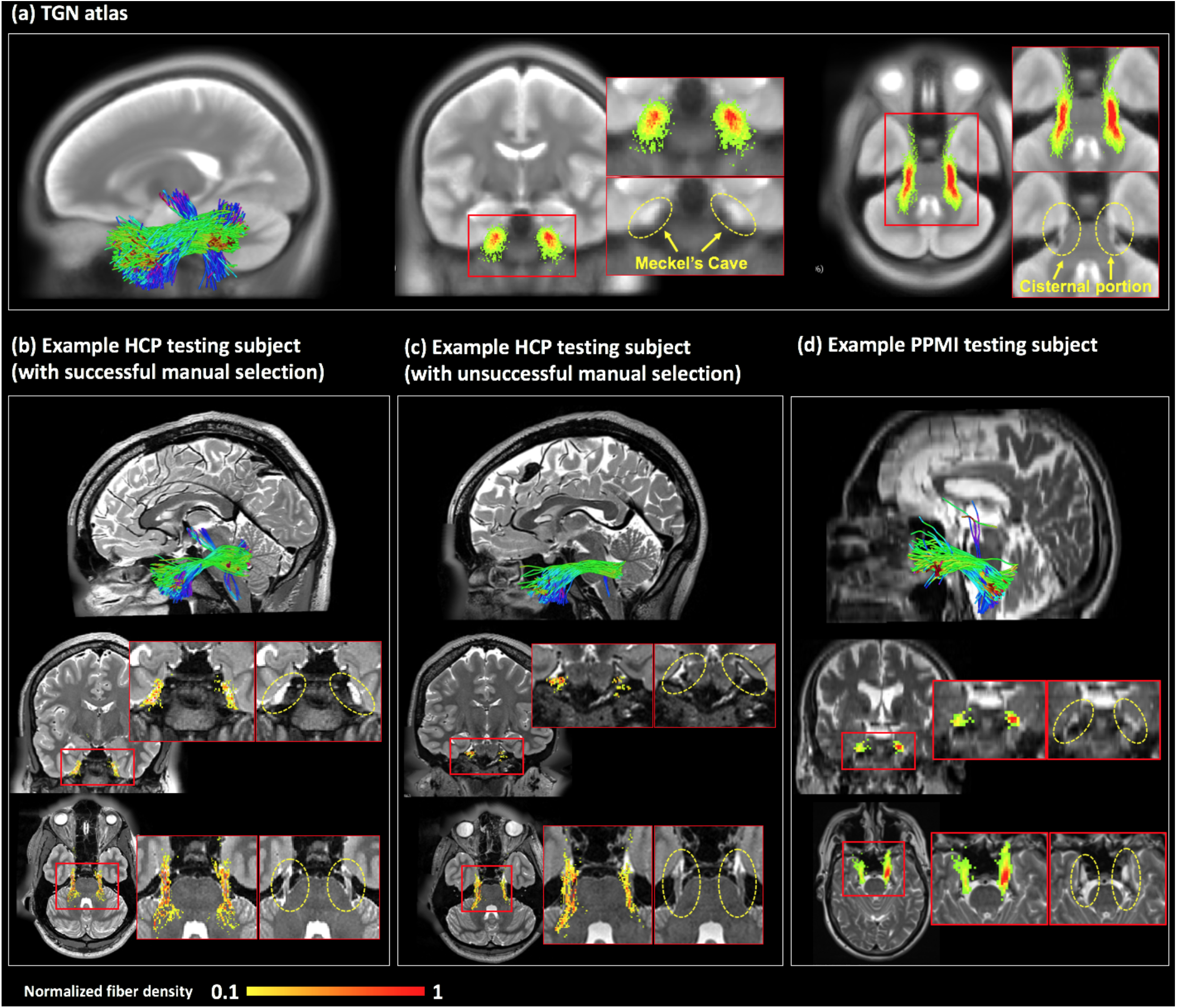
Visualization of the TGN 3D fiber trajectory and the voxel-based fiber density map, overlaid on T2w data. (a) The TGN in the atlas, overlaid on the population mean T2w image. (b, c) Subject-specific TGNs of the two example HCP subjects, overlaid on the corresponding T2w images. (d) Subject-specific TGNs of the example PPMI subject (healthy control), overlaid on the corresponding T2w image. For each sub-figure, inset images are provided for better visualization of the regions where the TGN passes through. The value of a voxel in the heatmaps represents the number of fibers that have fibers passing through the voxel. For visualization of the fiber density map at the same scale, each map is normalized by the maximal value on the map for each sub-figure.

## 4. Discussion

In this paper, we present a TGN tractography fiber clustering atlas to enable automated identification of TGN in dMRI tractography from new subjects. We show not only highly comparable TGN identification performance of our method with comparison to expert TGN identification, but also several advantages. First, our method performed automated TGN identification, without required expert ROI placement; thus, it provides an efficient tool to reduce expert labor costs and inter-operator bias. Second, our method was robust to potential imaging artifacts and/or noise and thus obtained a higher successful TGN identification rate. We have several overall observations about the results, which are discussed below.

We demonstrated successful application of the TGN atlas for subject-specific TGN identification, where 100% of the TGNs of all subjects under study were successfully identified. Importantly, our method could successfully identify the TGNs of the 8 HCP testing subjects where manual TGN selection could not because of failed ROI placement within the MC and the CP. In our work, we found that manual ROI placement was affected by imaging artifacts and/or noise at the skull base region from two aspects. First, ROIs could not be drawn because the anatomical structures of interest were not visible on the noisy dMRI data. Second, ROIs from inter-modality imaging (e.g. anantomcal T2w) could not be applied because of bad image registration at the skull base region. Our method identified the TGNs from dMRI tractography directly, without relying on the success of ROI placement. Therefore, our method provided a robust tool for TGN identification, in spite of the potential imaging artifacts and/or noise at the skull base region.

We showed the proposed atlas’s high TGN identification performance despite the heterogeneity of the dMRI data, specifically in the dMRI with relatively low spatial resolution. Our testing data included the dMRI data from the PPMI database, which was independently acquired using a different scanning protocol and processed in a different manner compared to the HCP data. These factors could affect the tractography results and thus influence the identification generalizations between different dMRI datasets. However, despite any potential effects from the heterogeneity of the dMRI data, we showed excellent TGN identification generalization performance across the multiple testing datasets. One important contributing factor was the application of the two-tensor UKF tractography (Malcolm et al., 2010; Reddy & Rathi, 2016) which is highly sensitive and robust in fiber tracking in dMRI data from different acquisition protocols.

We demonstrated the anatomical validity of the identified TGN using the proposed atlas. First, the automatically identified TGNs were highly comparable to the ground truth manual TGN selection results, where we showed highly visually similar TGN fiber trajectory and good spatial overlap. Second, the automatically identified TGNs corresponded well to the known anatomy, passing through the MC and overlapping with the CP, as appearing on the T2w image. T2w data has a good contrast of the cisternal portion of the TGN and has been widely used to confirm the presence of the TGN (Casselman et al., 2008; Xie et al., 2020).

The proposed atlas enabled identification of subdivisions of the TGNs. The TGN covers an extensive nerve distribution territory, including several segments such as the cisternal portion, the branching structures, the mesencephalic trigeminal tract, and the spinal trigeminal tract (Go et al., 2001; Joo et al., 2014). Unlike the cisternal portion of TGN that has been studied in multiple previous works, the mesencephalic trigeminal tract and spinal trigeminal tract are relatively less studied. The spinal trigeminal tract is important for mapping the pain-temperature sensory functions of the face, mouth and nose (Grant & Arvidsson, 1975). The mesencephalic trigeminal tract is an important portion of the TGN that conveys proprioceptive information from the teeth, masticatory muscles and temporomandibular joints (Shigenaga et al., 1989). To our knowledge, our recently published work (Xie et al., 2020) demonstrated, for the first time, the possibility of identification of the putative mesencephalic trigeminal tract using dMRI tractography techniques, where we have shown that the highly sensitive fiber tracking UKF algorithm can effectively track through the intra-brainstem region where the mesencephalic trigeminal tract fibers cross white matter fibers. In the present study, using the same underlying tractography method, we have shown a better identification rate using the proposed automated atlas-based method than previously applied manual selection method. However, while we showed modestly successful performance on identifying the mesencephalic trigeminal tract and spinal trigeminal tract, we noticed that the identification of these substructures could be affected by the image quality. In the PPMI data, we found a lower identification rate of these two tracts compared to the high-quality and high-resolution HCP data. This result suggested that improving the imaging acquisition would be helpful for identification of the more comprehensive anatomy of the TGN.

The proposed atlas-based TGN identification method aimed to address the known tractography issues of false negative and false positive fiber tracking (Maier-Hein et al., 2017; Thomas et al., 2014). In our study, we applied the multi-tensor UKF tractography method that has been shown to be highly sensitive tracking in the presence of crossing fibers and peritumoral edema in the cerebrum (Z. Chen et al., 2015, 2016; Gong et al., 2018; Hong et al., 2018; Liao et al., 2017; O’Donnell et al., 2017; F. Zhang, Kahali, et al., 2017; Fan Zhang, Savadjiev, et al., 2018; Fan Zhang, Wu, et al., 2018). The multi-tensor UKF tractography has also been demonstrated to be highly sensitive in tracking the different anatomical subdivisions of the TGN (Xie et al., 2020). The high sensitivity has been suggested to be important to reduce false negatives, but at the expense of increased false positive fiber tracking (Maier-Hein et al., 2017; Thomas et al., 2014). Therefore, the multi-tensor UKF fiber tracking method may introduce more false positive or anatomically incorrect errors compared to a standard single-fiber diffusion tensor fiber tracking method. In our method, we included two solutions to remove possible false positive fibers. First, during expert judgment, we excluded the fiber clusters that were anatomically incorrect to belong to the TGN. For instance, we found and excluded false positive fiber clusters entering the temporal lobe. While the expert judgement could reject the entire cluster if the cluster was not anatomically correct, improbable fibers within a cluster could not be ameliorated. To handle such tractography errors, we included a data-driven outlier fiber removal process to reject improbable fibers within a cluster.

While the aforementioned two processing steps have largely ameliorated the issue of false positive fibers, we still found false positive fibers entering the cerebellar peduncles, e.g., fibers that are parallel to the middle cerebellar peduncle. (See Supplementary Figure 4 for a graphic illustration of the false positive fibers entering the cerebellar peduncles). False positive tracking of the TGN has been reported in several studies, in particular, false positive streamlines entering the cerebellar peduncles (Behan et al., 2017; David Q. Chen, DeSouza, et al., 2016; David Qixiang Chen et al., 2011; Hung et al., 2017; Timothée Jacquesson et al., 2018; M. Yoshino et al., 2016). In our previous study that compared different fiber tracking strategies (dMRI data with different b-values, in combination with both single- and multi-tensor tractography methods), we have also found that false positive tracking into the cerebellar peduncle was a large challenge, as this false positive tracking was present in all datasets even with expert selection of TGN fibers (Xie et al., 2020). While this issue can be ameliorated by removing the fiber clusters that have fibers entering the cerebellar peduncles, unfortunately this strategy will remove a large number of fibers, reducing the possibility of identifying other structures such as the cisternal portion and the branching structures. Therefore, we included these fiber clusters in the curated TGN atlas. We note that there is a false negative tracking of the fibers that travel towards the trigeminal sensory nucleus, in particular the chief sensory nucleus; however, the TGN fibers to the other parts of the trigeminal sensory nucleus including the spinal nucleus and the mesencephalic nucleus can be identified using our method. (The trigeminal sensory nucleus is composed of three nuclei including the chief sensory nucleus, the spinal nucleus and the mesencephalic nucleus (Go et al., 2001; Joo et al., 2014).)

The proposed TGN atlas can be useful in both scientific and clinical applications. For example, our method provides a useful tool to enable large-scale population-wise statistical analysis. Many research studies have suggested that TGNs are important for understanding and/or potential treatment of various neurological disorders such as major depressive disorder, attention-deficit/hyperactivity disorder and Parkinson’s disease (Barz et al., 1997; McGough et al., 2015; Schrader et al., 2011). To identify disease-population-specific characteristics of the TGNs compared to healthy controls, statistical group-wise comparison is needed. Our automated method is efficient and can ensure a highly reliable population-wise statistical analysis, where TGN identification is performed in a consistent way across the subjects. Another example application of our TGN atlas is to identify and locate the vulnerable TGNs in tumor patients for surgical planning research. In particular, presurgical visualization of TGN displacement due to tumor/lesion compressions offers a significant asset to predict the vulnerability of the TGNs in neurosurgery (Timothée Jacquesson et al., 2018). Our atlas can also be useful by providing a possibility to quantify the spatial CN trajectory deviation (displacement due to peritumoral effects) from the TGNs of healthy brains in the atlas. Another potential use of our method is to study TGN pathologies related to neurovascular conflict, e.g., in trigeminal neuralgia. Our method uses diffusion MRI tractography, which is sensitive to water diffusion and fiber myelination but not vessels or arteries. In combination with other MRI modalities where vessels and arteries are highly visualized (Donahue et al., 2017; Haller et al., 2016; Kontzialis & Kocak, 2017), our method can provide an effective tool to separately identify nerve structures to confirm neurovascular conflict.

Potential limitations of the present study, including suggested future work to address limitations, are as follows. First, in the present study, we demonstrated improvement to the manual ROI-based TGN selection methods on the testing HCP datasets that were affected by imaging artifacts and/or noise. To perform consistent processing across subjects and provide fairly comparable results to the literature, we chose a widely used method based on predefined manually drawn ROIs within the MC and CP. For this particular manual selection strategy, ROI placement was affected by imaging artifacts and/or noise so that TGN selection was not able to be performed. However, we acknowledge that a manual TGN identification method that requires more sophisticated processing, e.g., interactively moving ROIs (Chamberland et al., 2012; Golby et al., 2011; Fan Zhang et al., 2020), may also identify the TGNs in these testing datasets (see Supplementary Figure 3). Second, we demonstrated our method on TGN identification of subjects with different health conditions, including healthy and Parkinson’s disease. A further evaluation could include an investigation of patients with secondary pathologies that affect the TGN, e.g. trigeminal neuralgia and neurosurgical patients with skull base tumors. Third, in this study, we created the TGN atlas using UKF tractography because it has been demonstrated to be effective in tracking TGNs (Xie et al., 2020). In an initial experiment, we have shown successful applications of the atlas to tractography data computed using two additional fiber tracking methods, including diffusion tensor tractography (Basser et al., 2000) and constrained spherical deconvolution (Jeurissen et al., 2011) tractography (see Supplementary Material S1 for details). An interesting future work could include a comprehensive comparison to investigate the differences of the TGNs identified from different tractography methods. Fourth, given the success in atlas curation of the TGN, as well as the brain white matter (Fan Zhang, Wu, et al., 2018), we believe that it is highly feasible and promising to leverage our fiber clustering techniques for atlas curation of other cranial nerves, which is interesting further work to be investigated. Fifth, a more comprehensive assessment of the automatically identified TGN fibers could include a comparison to advanced CISS or FIESTA data that can provide better visualization of the cisternal portion of the TGN. However, due to the unavailability of such advanced data in the HCP and PPMI datasets under study, we chose to use the T2w data that provide reasonably good contrast of the cisternal portion of the TGN and have been widely used to confirm the presence of the TGN (Casselman et al., 2008; Xie et al., 2020).

## 5. Conclusions

In this paper, we have presented a novel dMRI tractography fiber clustering atlas that enables automated identification of the TGN of new subjects. Experimental results show successful application of the proposed atlas to dMRI data with different MRI acquisition protocols and demonstrate advantages over a traditional manual selection strategy.

## ACKNOWLEDGMENTS

We gratefully acknowledge funding provided by the Jennifer Oppenheimer Cancer Research Initiative and the following National Institutes of Health (NIH) grants: P41 EB015902, P41 EB015898, R01 MH074794, R01 MH111917, R01 MH119222, R01 CA235589, HHSN261200800001E and U01 CA199459.

## Footnotes

1 https://github.com/SlicerDMRI/whitematteranalysis

2 https://dmri.slicer.org/atlases

3 https://dmri.slicer.org/

4 https://github.com/pnlbwh/ukftractography

5 https://github.com/SlicerDMRI/whitematteranalysis

