## supplementary materials for "Creation of a novel trigeminal tractography atlas for automated trigeminal nerve identification"

**S1. Application of the TGN atlas to tractography computed from different fiber tracking methods**

In this supplementary material, we provide the results from applying the proposed TGN atlas to the tractography data generated using two additional fiber tracking methods (different from the UKF method that was used for the atlas generation). The two additional tractography methods included a traditional second-order Runge-Kutta single-tensor DTI streamline method (Basser et al. 2000) (implemented in 3D Slicer via SlicerDMRI (Norton et al. 2017; Zhang et al. 2020)) and an advanced second-order integration over fiber orientation distributions (iFOD2) CSD method (J. D. Tournier, Calamante, and Connelly 2010) (implemented in MRtrix3 (J.-D. Tournier et al. 2019)). Data from one example HCP subject was used in this experiment.

Parameters for the two methods were set as follows. For the DTI method, tractography was seeded within the TGN tractography seeding mask in all voxels where FA was greater than 0.06, and stopped when FA fell below 0.05. For the CSD method, the fiber orientation distribution (FOD) was computed (Tournier et al., 2007), and fiber tracking was performed using an iFOD2 method (Tournier et al., 2010) with the default parameters as suggested by the software. For each of these two methods, around 30,000 fibers were computed. We also give the results using the UKF tractography for comparison. For the UKF method, we used the parameters as reported in the main paper content (Section 2.1.2), except for that we used *5 seeds per voxel* to increase fiber density, so that the number of fibers in the tractography data (about 30,000) was similar to those from the DTI and CSD methods.

Figure S1 gives a visualization of the TGNs obtained using the three different tractography methods, including the DTI, the CSD, and the UKF methods. In general, the obtained tracts were visually plausible, showing the ability of the proposed atlas in generalizing to tractography data computed using different tractography methods. However, given that the tractography data varied quite significantly across the fiber tracking models, we also observe several differences. First, the DTI method could hardly track the fibers belonging to the putative mesencephalic trigeminal tract, while the CSD and UKF methods managed to track these fibers. Second, the UKF method is more sensitive in tracking the branching structures than the other two methods. Third, the UKF and DTI tractography are deterministic fiber tracking methods, generating fiber streamlines that are relatively smooth compared to the CSD probabilistic method.

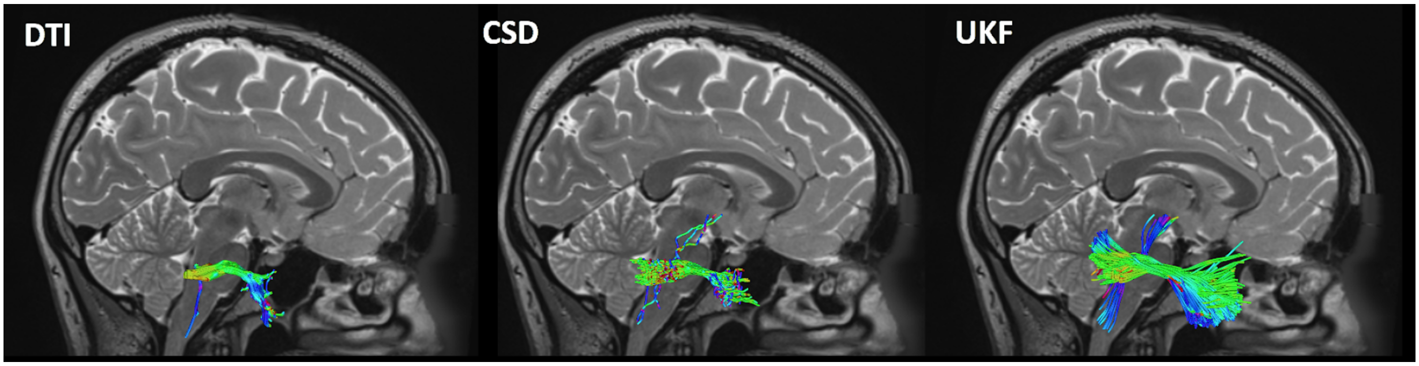


Figure S1. Visualizations of the TGNs identified using three different tractography methods: the UKF method (left), the CSD method (middle), and the DTI method (right).

**S2. Supplementary Figures:**

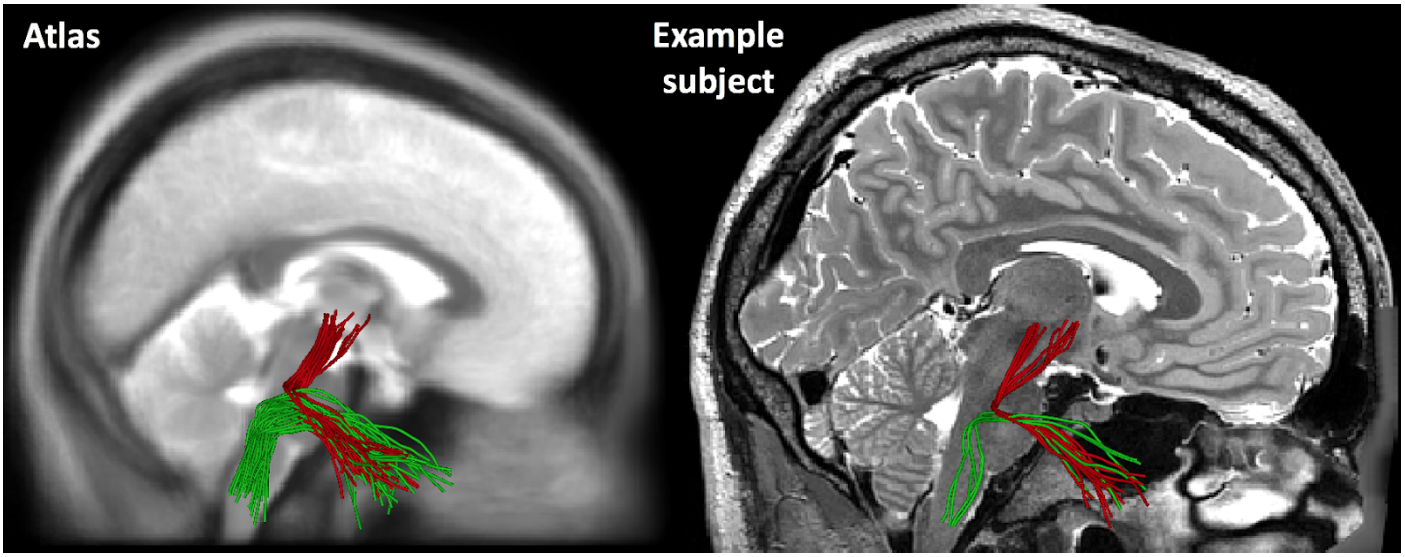


Supplementary Figure 2. A visualization of the intra-brainstem portion of the TGN in the curated tractography atlas and the corresponding structures identified in one example HCP subject.

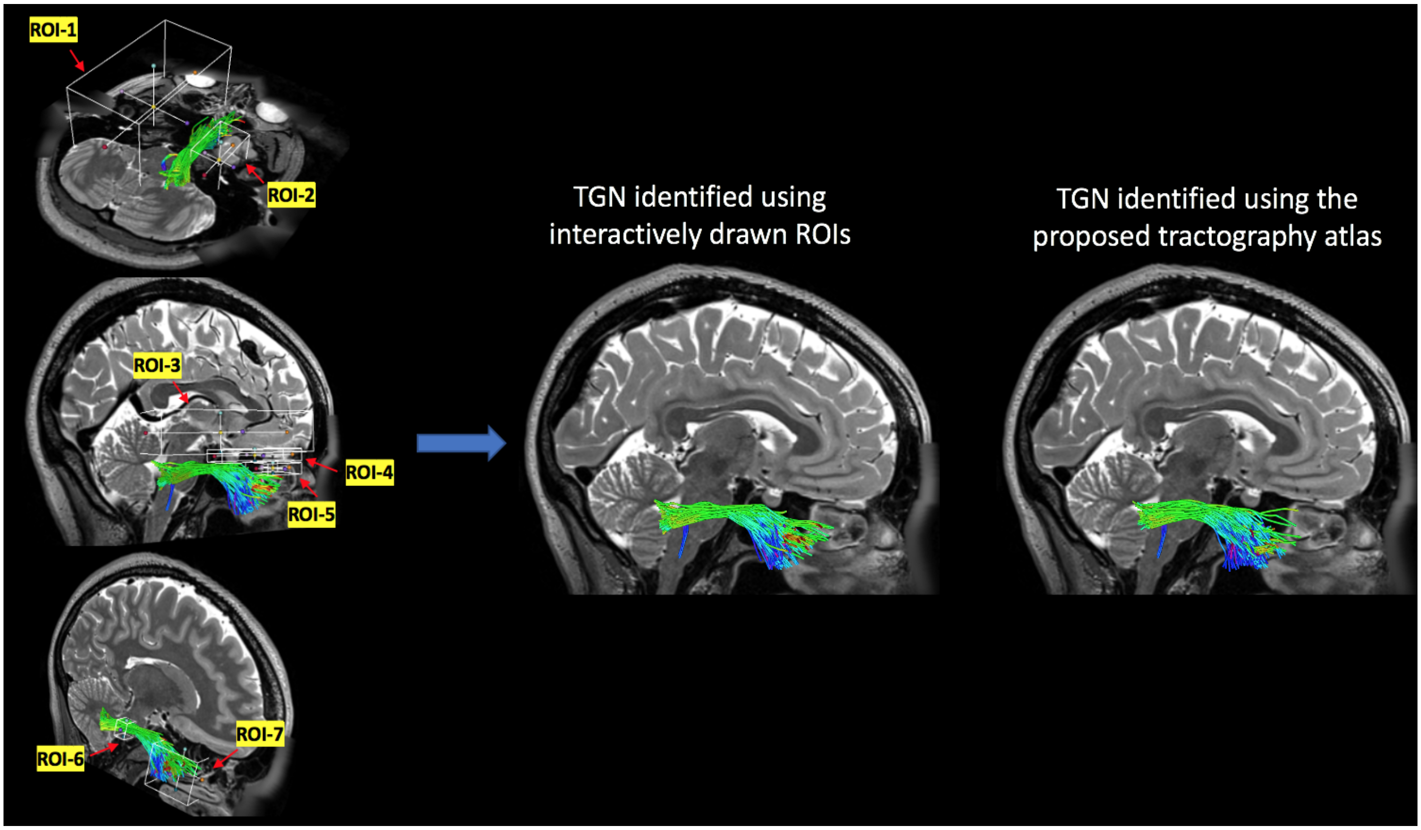


Supplementary Figure 3. TGN identified using interactively drawn ROIs on one example HCP dataset (the same subject as in Figure 3b in the main paper), where TGN selection using the predefined ROIs failed. This interactive ROI TGN selection was performed by an expert (FZ), who has been trained with neuroanatomical knowledge of the TGN. The "Fiber Bundle Section" functionality within the "Tractography Display" module in the SlicerDMRI software was used, which allows for interactively placing the ROIs, adjusting their sizes, and viewing fiber selection results. 7 ROIs were used with the following purposes: ROI-1 for exclusion of fibers in the contralateral side of the skull base; ROI-2 for exclusion of fibers entering the temporal lobe; ROIs-3,-4 and

-5 for exlusion of fibers entering the frontal lobe; ROI-6 for inclusion of fibers passing through the cisternal portion of the TGN as appearing on the T2w data (note this ROI is larger than the actual size of the cisternal portion in order to account for small registration errors between the T2w and dMRI data); ROI-7 for including of fibers belonging to the TGN branching structures. The identified TGN using these interactively drawn ROIs has an anatomically correct shape, corresponding to the known anatomy of the TGN pathways. The automatically identified TGN using the proposed tractography atlas has a visually comparable identified result compared to the interactively selected TGN.

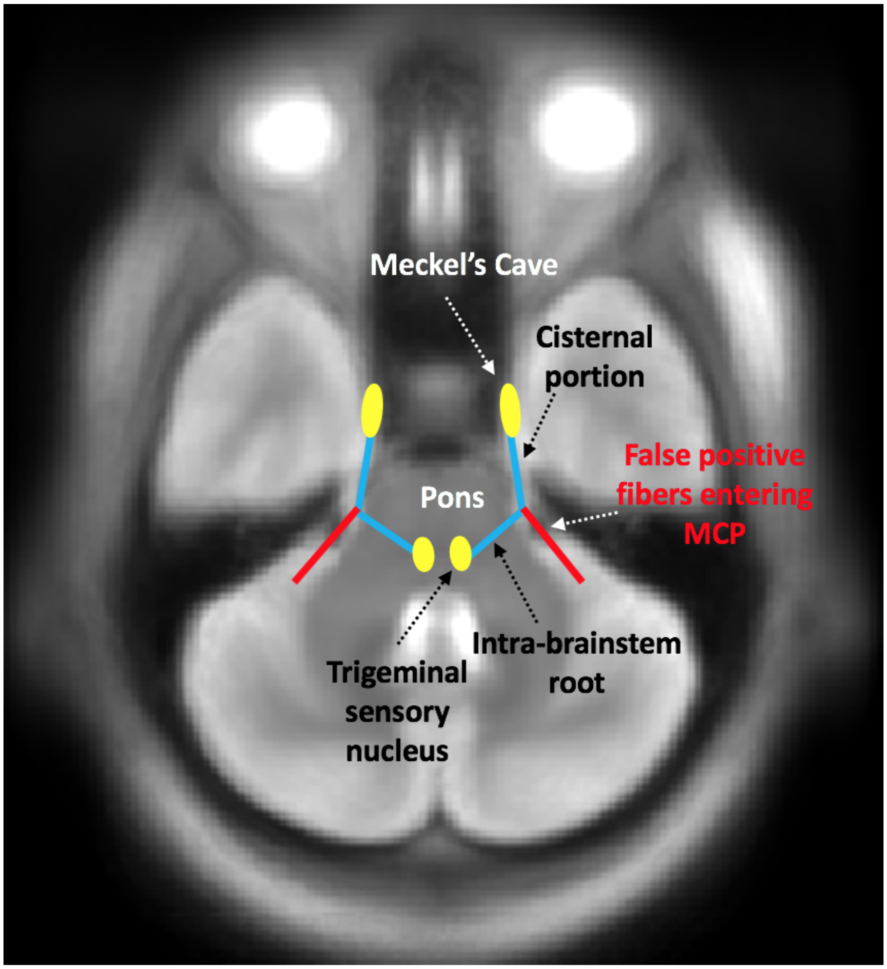


Supplementary Figure 4. A graphic illustration of the false-positive TGN tracking entering the middle cerebellar peduncle (MCP).
